# Withaferin A downregulates NDRG1 to overcome hypoxia mediated EMT and chemoresistance in lung adenocarcinoma and glioma cells

**DOI:** 10.1101/2024.12.26.629226

**Authors:** Divyank Mahajan, Sanchit Gandhi, Tapasya Srivastava

## Abstract

One of the important hurdles to be overcome in the management of cancers is the acquired resistance to therapy. There is a crucial need to not only understand molecular pathways underpinning resistance to therapy but to find better and more effective therapeutic avenues to overcome such acquired resistance. Tumor microenvironmental conditions such as hypoxia have long been known to result in chemoresistance in various cancers that become unresponsive to standard-of-care drugs such as cisplatin. Although many new compounds are screened for their anticancer properties, the recalcitrant tumor microenviroenment has not been given due importance in many potential drugs. In this study, we examine the efficacy of Withaferin A (WA), a bioactive compound isolated from *Withania somnifera*, in overcoming hypoxia-induced metabolic adaptation, chemoresistance, and epithelial-to-mesenchymal transition (EMT) in both lung adenocarcinoma and glioma cells. Our results reveal that WA significantly inhibits hypoxia-induced increased migration, EMT, and metabolic reprogramming, leading to enhanced cytotoxicity in otherwise chemoresistant cancer cells. WA treatment suppressed the expression of markers of EMT, glucose metabolism, and bonafide hypoxia markers, particularly NDRG1 thereby inducing cell death in these cancer cells. WA showed significant cytotoxic effects in chemoresistant lung adenocarcinoma and glioma cells, both alone or in combination with cisplatin, highlighting its potential as a therapeutic agent for overcoming chemoresistance in these cancers. Our results provide new insights into the anti-cancer mechanisms of WA especially under hypoxic conditions, and support its further investigation as a promising adjunctive therapy for the treatment against hypoxia-induced chemoresistance in cancers.

## Introduction

Tumor hypoxia has been implicated as an important player in cancer progression and metastasis. Due to the relentless proliferation of tumor cells, a tumor exceeds the size greater than the diffusion capacity of oxygen from the blood vessels present at its periphery, which creates a limiting supply of oxygen (and nutrients) for the cells located spatially inside of the tumor. Under such conditions of limiting oxygen termed tumor hypoxia, cancer cells globally re-adapt cellular machinery to maintain their survival and proliferation. As part of the cancer cells make either imperfect attempts to form new blood capillaries or attempt a change in the tone of existing capillaries i.e. dilation of blood capillaries and vessels. These re-adaptation strategies manifest as an increase in metastatic potential, aggressiveness, invasiveness, and increased resistance towards chemotherapy and radiotherapy. In addition, hypoxic environments also potentiate the generation of cancer stem cell [1]. Hypoxia, therefore, has emerged to be a major concern in tumor pathophysiology since it advances tumor growth and resistance against therapies in a wide range of malignancies such as cancers of the lung, breast, prostate, brain, uterine, vulva, head and neck [2].

Co-opting fundamental cellular signalling pathways, hypoxia contributes to the ‘hallmarks of cancer’ - sustained proliferative activity, evading growth suppressors, activating invasion and metastasis, enabling replicative immortality, inducing angiogenesis and resisting cell death [3]. HIF-1α and HIF-2α are observed to be commonly upregulated in NSCLC in a correlated manner, and are also correlated with poor prognosis; HIF-2α is more strongly correlated with poor prognosis [4].

Withaferin A (WA), a biologically active steroidal lactone extracted from *Withania somnifera*, has long been known for its anti-cancerous properties. It has been shown to manifest anticancer properties through multiple mechanisms in various cancers of different tissue origins, including glioblastoma, neuroblastoma, lung, breast, colon, and head and neck cancers. It has been combined with different anti-cancer compounds in various cancer cell-lines to study improved efficacy. For e.g., WA has been studied in combination with TRAIL, Sorafenib, Doxorubicin, Etoposide, Celecoxib, Cisplatin, and Oxaliplatin in renal, papillary, and anaplastic, epithelial ovarian, breast, and pancreatic cancer cells, respectively [5–10]. In ovarian cancer cells, where tumor relapse is frequently observed as a consequence of platinum resistance, combining platinum-based drugs such as cisplatin with WA has been shown to exert a synergistic effect. The combination was shown to induce cell death by a generation of reactive oxygen species (ROS) generation and DNA damage requiring a cisplatin dosage lower than normal. WA is known to have a pleiotropic mechanism of molecular action. It has been shown that by binding to VEGF, WA inhibits angiogenesis [11].

The use of nude mice bearing orthotopic ovarian tumors also demonstrated WA’s capacity to eliminate putative cancer stem cells (CSCs) and the associated chemoresistance they exhibit [9,12]. Kyakulaga, AH et al. (2018), showed that treatment with Withaferin A of NSCLC cell lines -A549 & H1299-in a time and dose-dependent manner curbs proliferation and TGF-β induced EMT [13].

## Materials and Methods

### Cell lines, cell culture conditions

All cell lines used - A549, A172, T-98G were procured from National Centre for Cell Science (NCCS) cell line repository at Pune. All cell lines were cultured in Dulbecco’s Modified Eagle Medium (DMEM), Gibco, USA, supplemented with 10% Fetal Bovine Serum (Gibco, USA) at 37°C and 5% CO2 under >95% relative humidity. The growing cells were replenished with fresh medium every 2-3 days. Cells were passaged at 70-80% confluency by washing the monolayer with DPBS and harvesting using trypsin-EDTA solution. To expose cells to hypoxic conditions, cells were kept in an incubator chamber Anoxomat chambers (Mart® Microbiology and the Anoxomat™ system) and infused with a regulated gas mixture of 5% CO2, 1%O2 and 94% N2 and incubated at 37°C.

### Drug treatments

#### Withaferin A preparation

Withaferin A was purchased from Sigma Aldrich (Cat No: 681535), CAS No-5119-48-2-Calbiochem. A stock solution of 2 mg/mL was prepared by dissolving the crystalline white powder in absolute ethanol and aliquots were stored at -20°C.

#### Cisplatin treatment

Cisplatin was purchased from Sigma Aldrich (Cat No: 232120) (CAS No-15663-27-1-Calbiochem). A stock of 1 mg/mL was prepared by dissolving the crystalline white powder in PBS.

### Fluorescence Microscopy

A549 cells were treated with three different concentrations (0, 3 and 6 μM) of WA in normoxia for 8 hours and F-actin making up the cytoskeleton were stained using Phalloidin. The cells were counterstained with DAPI for identifying the nuclei. Cells were visualized and photographed on a confocal microscope.

### RNA Isolation & Real-time PCR

A549 cells were cultured in 60mm dishes or in 6-well plates. After the cells had attained a confluence of 70-80%, experimental perturbations (treatment or hypoxia exposure) were made, and cells were incubated in respective growth conditions for a designated time period. Total RNA was isolated from the cells exposed using the Trizol method (Thermo Fisher Scientific). RNA pellet was dissolved in 20*µ*L RNase-free water and incubated at 55°C for 10 minutes to allow for a complete dissolution of RNA, and assessed using Nanodrop (Thermo Scientific, USA). First-strand cDNA synthesis was done using RevertAid RT Reverse Transcription Kit (Thermo Fisher Scientific) with the random hexamers according to the manufacturer’s protocol using 1µg of total RNA from each sample.

Primer for each gene was either manually designed or using the NCBI primer tool and checked for specificity using primer BLAST (Supplementary Table 1). qPCR reaction mixtures were of 10µl and contained Syto9 as the intercalating dye. qPCR reactions were performed on Qiagen’s Rotor-Gene-Q instrument and analyzed using Rotor-gene Q series software-2.3.1.49. Cycle threshold (Ct) values were normalized to amplification of the 18S transcript and the data were analyzed using the 2(-Delta Delta Ct) method by Livak and Schmittgen [14].

### Wound healing assay (Scratch-assay)

A monolayer of cells was allowed to grow to a confluency of 70% in a well of a 6-well plate, post which a vertical wound was made across the monolayer using a 200µL pipette tip. Growth medium containing either WA or vehicle at the appropriate concentrations was added. Images were captured at regular intervals as per the design of the experiment.

### Cell viability assay & MTT Assay

1×10^4^ cells per well were seeded in 96-well plates and cultured overnight and were treated with various concentrations of WA. Post-treatment, after removing the spent media, MTT reagent (200µg/mL) was added to each well and incubated at 37°C for 3 hours in the dark and 200μL DMSO was added to dissolve formazan crystals. The number of viable cells was assessed by determining absorbance measured at 570 nm in a plate reader (TECAN, Switzerland). Cell viability percent was calculated as Ab (treatment)/Ab (vehicle) x 100. (MTT stock solution was prepared by dissolving 2 mg MTT in 1 ml of PBS. The solution was filtered through a 0.22 μm pore filter and aliquots were stored at -20°C).

### Protein lysate preparation and western blotting

Cells having reached a confluence of 70-80% were exposed to hypoxia or treated with WA and exposed to hypoxia as per the experiment design and requirements. After the indicated time of the experiment, cell culture dishes were immediately kept on ice. Cells were rinsed twice using ice-cold PBS solution and lysate was prepared using 1X RIPA lysis buffer and stored at -80°C.

An equal amount of each sample was loaded in respective wells. Gel was then run at a constant voltage of 100-120 V for 1 hour. Transfer was performed at 4°C overnight at a constant voltage of 40V. All membranes were washed with TBS and blocked with 5% skimmed milk in TBST for 2 hours. Post-blocking, membranes were incubated with primary antibodies prepared at an appropriate solution.

### Expression Analysis using GEPIA

The GEPIA database (http://gepia.cancer-pku.cn/index.html) was used to compare expression levels of selected genes in GBM and LUAD with their matched normal tissues. It utilizes public RNA sequencing data from The Cancer Genome Atlas TCGA and the Genome Tissue Expression (GTEx) database to perform differential gene expression analysis. Here, expression data were log2-transformed as per log_2_(TPM+1) while log2FC was defined as median tumor – median normal. p-value threshold was kept at 0.01.

## Results

### 1. Withaferin A reduces EMT and downregulates molecular markers of metastasis, hypoxia and apoptosis

We observed that in response to treatment with WA, expression of bonafide markers of hypoxia - Glut-1 and Pdk-1 are downregulated. A549 cells were treated with WA for 24 hrs at two concentrations 1µM and 4µM in normoxia and 1% hypoxia. qPCR analysis revealed that hypoxia exposure upregulated expression of both Glut-1 and Pdk-1 more than 5 folds relative to normoxia (Fig 1). Cell survival or death depends upon the balance between pro-apoptotic signals or anti-apoptotic signals [15,16]. We also observed changes in pro-apoptotic markers – p53, Bax, and p21, after 24 hours of WA treatment. WA treatment upregulates the expression of pro-apoptotic Bax, particularly at 4µM concentration in hypoxia. In normoxia, Bax was observed to be upregulated at low dose, p21 was largely unchanged and p53 showed upregulation at high dose of Withaferin. In normoxia, a dose dependent trend was not seen. Overall, pro-apoptotic genes appeared to be enhanced (Supplementary figure 1)

**Figure 1(A-D):**
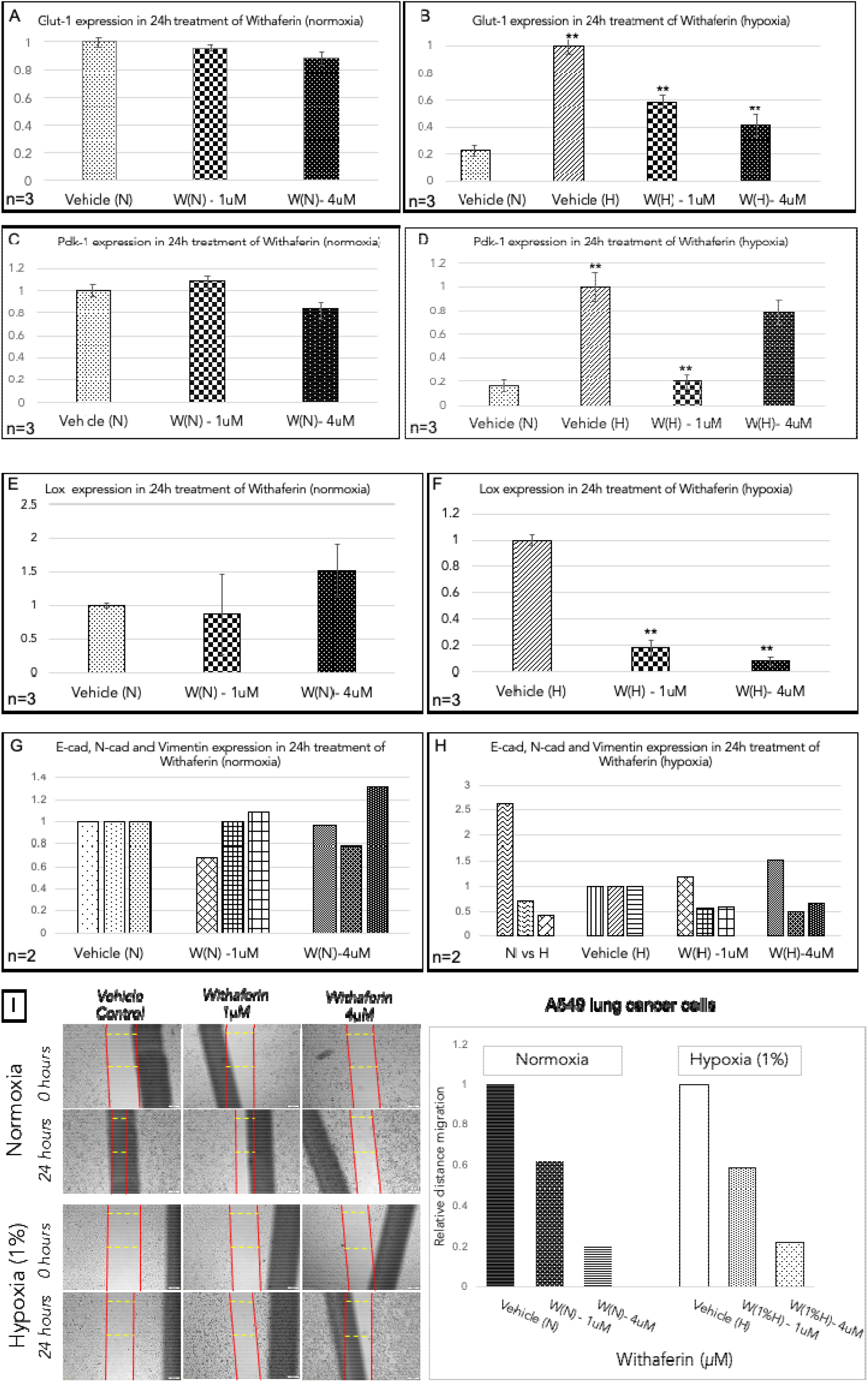
Gene expression of hypoxia markers in A459. (A) Glut-1 expression is not significantly changed with 24h treatment of WA at 1µM and 4µM wrt control (vehicle). (B) Glut-1 expression is significantly higher in hypoxia (vehicle) wrt to normoxia (vehicle control) (p<0.01). For the benefit of depiction, the values have been normalized to hypoxia vehicle control. Glut-1 is significantly downregulated with Withaferin treatment in hypoxia conditions at both doses (p<0.05). (C) Pdk-1 expression is not significantly changed with 24h treatment of WA at 1µM and 4µM w.r.t control (vehicle). (D) Pdk-1 expression is significantly more in hypoxia (vehicle) w.r.t to normoxia (vehicle control) (p<0.01) and is significantly downregulated with Withaferin treatment in hypoxia conditions at 1µM WA (p<0.05). For the benefit of depiction, the values have been normalized to hypoxia vehicle control. Y axis depicts fold change values with respect to Vehicle (N) in A and C and Vehicle (H) in B and D. **(E-I)**: Gene expression of EMT, invasion and metastasis markers in A459 (E) Lox expression is not significantly changed with 24h treatment of WA at 1µM and increased in 4µM w.r.t control (vehicle) although not significant. (F) Lox expression is significantly reduced with treatment of Withaferin in hypoxia w.r.t to hypoxia (vehicle control) (p<0.01) in both doses. (G) E-cadherin, N-cadherin and Vimentin expression were studied after Withaferin treatment in normoxia (n=2). E-cadherin was reduced in 1µM Withaferin while N-cadherin was reduced in 4µM withaferin. Vimentin was increased in both doses. (H) E-cadherin, N-cadherin and Vimentin expression were studied after Withaferin treatment in hypoxia. E-cadherin expression is higher in Withaferin treated cells w.r.t to hypoxia (vehicle control) while both N-Cadherin and Vimentin expression were reduced with WA treatment compared with hypoxia (vehicle) Y axis depicts fold change values with respect to Vehicle (N) in E and G and Vehicle (H) in F and H. (I) Wound healing assay was performed to assess the change in migration potential. There was a dose dependent decrease in distance migration on withaferin treatment in both normoxia and hypoxia

It is established that readaptation strategies in response to tumor hypoxia initiate and promote metastasis, and consequently, is associated with poor survival in a variety of solid tumors [17,18]. Stabilized HIFs transactivate a number of players such as L1 cell adhesion molecule, factors of epithelial to mesenchymal transition (EMT), and the lysyl oxidase family members-LOX, LOXL2, and LOXL4 which promote extravasation and play crucial roles in invasion and formation of metastatic niches [19,20]. LOX has been proposed to be a promising target to treat progressive and metastatic cancer [21]. In order to gain insights into the effect of WA treatment upon the metastatic potential of A549 cells in hypoxia, we measured changes in the levels of known hypoxia-induced metastatic and EMT markers. Firstly, levels of LOX mRNA were evaluated via qPCR in response to WA treatment in A549 cells in normoxia and 1% hypoxia. WA treatment in hypoxia at concentrations of 1µM and 4µM downregulates expression of LOX by five and ten folds, respectively. Next, change in the level of EMT markers - E-cadherin, N-cadherin, and Vimentin, was evaluated in response to WA treatment in normoxia and hypoxia. In control samples, in response to hypoxia while the E-cadherin levels decrease, the levels of N-Cadherin and vimentin increase. When treated with WA, an increase in levels of E-cadherin expression was observed, and a significant decrease in levels of N-Cadherin and vimentin - which are promoters of mesenchymal states in cancer cells.

### 2. Withaferin A treatment disrupts the cytoskeleton and induces dose dependent cytotoxicity in lung cancer cells in both normoxia and hypoxia

Confocal microscopy for phalloidin after WA treatment showed disruption of the cytoskeleton in A549 cells in a dose dependent manner and the cytotoxic effect of WA treatment on cellular viability in hypoxic conditions vis-à-vis normoxic conditions was screened for at different concentrations ranging from (0.25µM to 6µM) in various cancer cell-lines including, A549. WA treatment (1µM) demonstrated a dose-dependent cytotoxicity in both normoxic as well as hypoxic (3% and 1%) conditions. WA treatment decreased viability to 63.49%, 62.05% and 63.29% in normoxia (p value = 0.028), 3% O_2_ (p value = 0.025) and 1% O_2_ (p value = 0.007) hypoxia respectively. We next pursued efficacy of a combination of WA at a lower dose with cisplatin, which is a standard-of-care drug used in solid tumors. This combination of WA (1µM) and cisplatin (5µg/ml) was used to treat A549 cells to ascertain the combined efficacy. The combination treatment resulted in a decreased viability to 54.21%, 39.94% and 55.25% in normoxia (p value = 0.02), 3% O_2_ (p value = 0.0006) and 1% O_2_ (p value = 0.002) hypoxia, respectively.

### 3. Withaferin A effectively increases cytotoxicity in chemo-resistant glioma cells and in liver cancer cells

Hypoxic tumour microenvironment plays an important role in brain tumour cells particularly gliomas. Our previous studies as well as others have determined that important role of hypoxia in determining the progression and drug responsiveness of glioma cells. Therefore, the effect of WA in the hypoxic microenvironment was also studied in A172 and T98G glioma cells. Treatment with just 1µM and 2µM WA for 48 hours and 72 hours was equally effective in A172 cells as 5µg/ml Cisplatin, the standard-of-care drug at its known physiological dose, in normoxia and hypoxia (Figure 2C). In T98G cells that are well-known to show resistance to the standard-of-care drug temozolomide (TMZ) (Figure D), WA appeared to be highly effective in a dose-dependent manner in all three conditions – normoxia, 0.2% and 1% hypoxia. Finally, WA was also effective in the liver cells, HEPG2 in both normoxia and hypoxia (Supplementary Figure 2).

**Figure 2:**
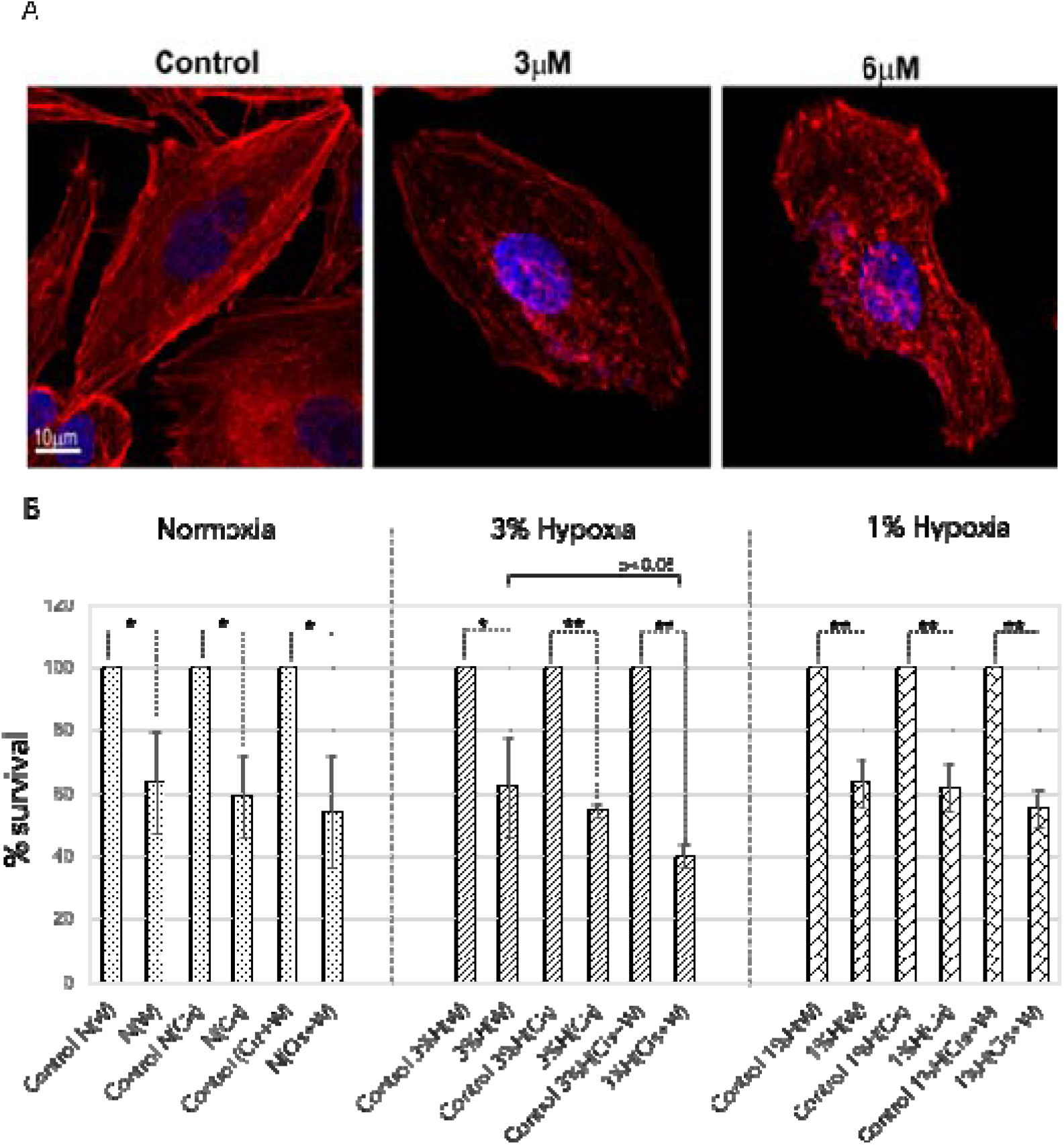
A) Representative microphotographs of A549 cells stained with phalloidin (Red) and DAPI (Blue) after treatment with WA (W) in normoxic (N) conditions for 8 hours showing stress fibers and cytoskeleton disruption. B) Combination of Cisplatin (Cis) and WA were evaluated for cytotoxicity in normoxia and hypoxia (3%H and 1%H). Cytotoxicity with WA alone was significantly higher in all groups. Combination of WA and Cisplatin was significantly more cytotoxic than WA alone in 3% hypoxia and as effective as WA and Cisplatin alone in other groups.

**Figure 3:**
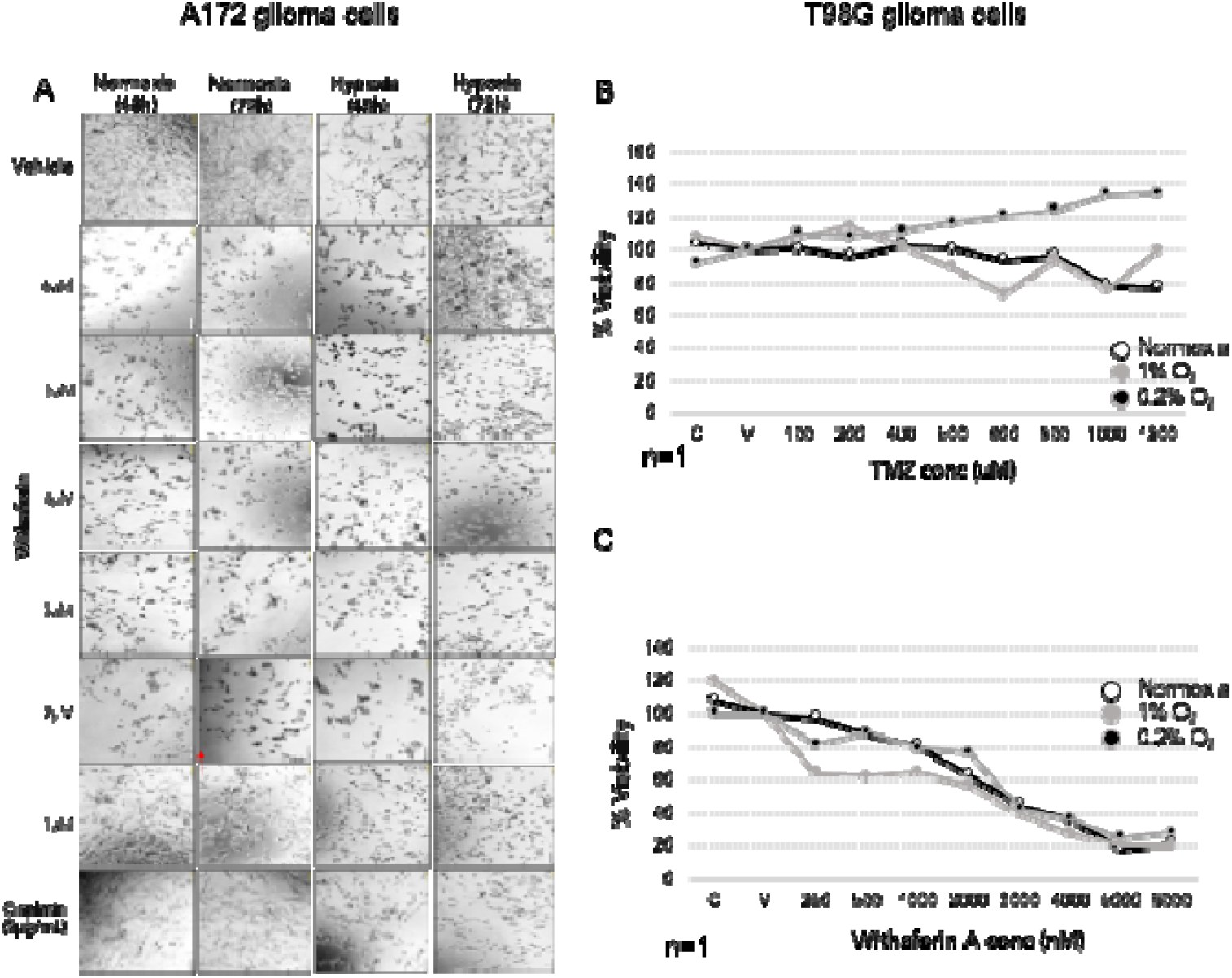
Withaferin induced cytotoxicity in A172 and T98G glioma cells. A) Representative microphotographs of A172 cells treated with WA for 48 and 72 hours in both normoxia and hypoxia show morphological changes and cytotoxic effect of 1µM and 2µM WA treatment as equivalent to the standard Cisplatin treatment (5µg/ml). B) T98G cells are known to be resistant to Temozolomide. Viability in normoxia and 0.1% and 1% O2 conditions on TMZ treatment shows poor cytotoxicity. C) Treatment of T98G cells with WA showed that these cells were much more responsive to WA and showed a dose dependent decrease in viability in all three conditions of heterogeneous oxygen depletion.

### 4. Withaferin A downregulates NDRG1, a bonafide marker of hypoxia

NDRG1 is a well-known marker for hypoxia and is often associated with poor outcome of many cancers. NDRG1 was determined to be highly upregulated (p value = 0.018 and 1.36×10^−59^, respectively) in analysis of two datasets (GSE200204 & GSE131378) of lung cancer cell-lines treated with hypoxia (Figure 4A). To evaluate the effect of the treatment of WA on various protein markers relevant to tumor hypoxia, immunoblotting was performed for N-Cadherin, NDRG1, and HIF-2α in both normoxic as well as 1% hypoxic conditions. It was observed that upon treatment with WA, expression of N-Cadherin, HIF-2α and NDRG1 decrease in a dose-dependent manner (Figure 4B). Downregulation of NDRG1 was evident in both 3% and 1% Hypoxia (Figure 4C) at 4µM WA indicating the effectiveness across variable hypoxia conditions which is a feature of solid tumour interiors. Clear down-regulation of NDRG-1 and HIF-2α, both essential for adaptation in tumor hypoxia, corroborate our findings and supports our hypothesis that treatment with WA helps overcome hypoxic advantage by downregulating key protein markers involved in adaptation and enhanced metastasis.

**Figure 4:**
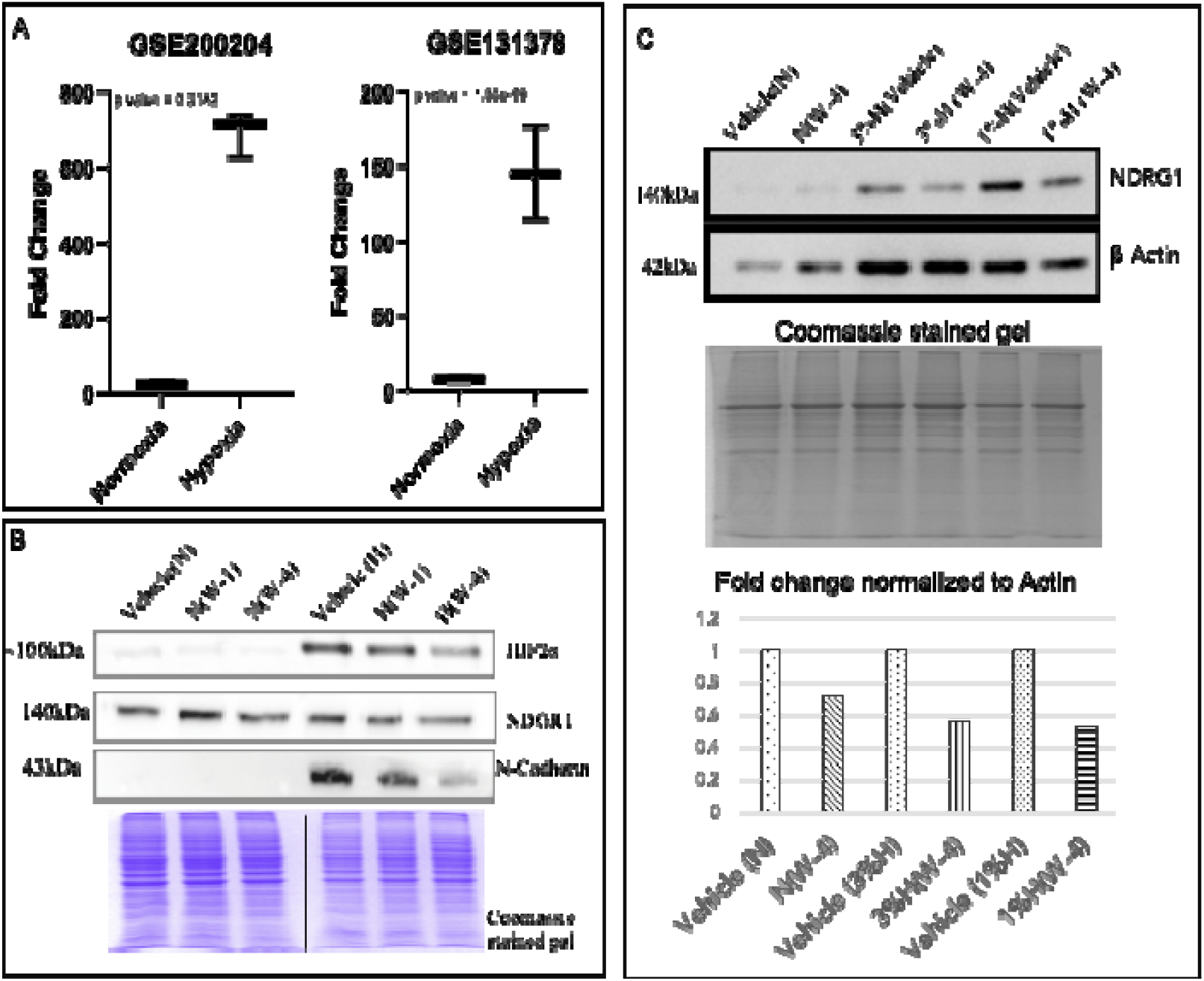
Effect of WA on hypoxia markers NDRG1 and HIF2A. (A) Analysis of two datasets of hypoxia cultured lung adenocarcinoma cells reveals a steep upregulation of NDRG1 in the RNA expression. (B) Treatment with WA at dose 1µM and 4µM downregulates HIF2A, NDRG1 and N-cadherin in 1% hypoxia. (C) Representative protein expression analysis - Western blotting, Coomassie stained gel for equal loading and fold change analysis is represented. WA downregulates NDRG1 in both 1% and 3% hypoxia conditions at 4µM. N: normoxia; H: hypoxia; W: WA, W-1: 1µM WA; W-4: 4µM WA.

**Figure 5:**
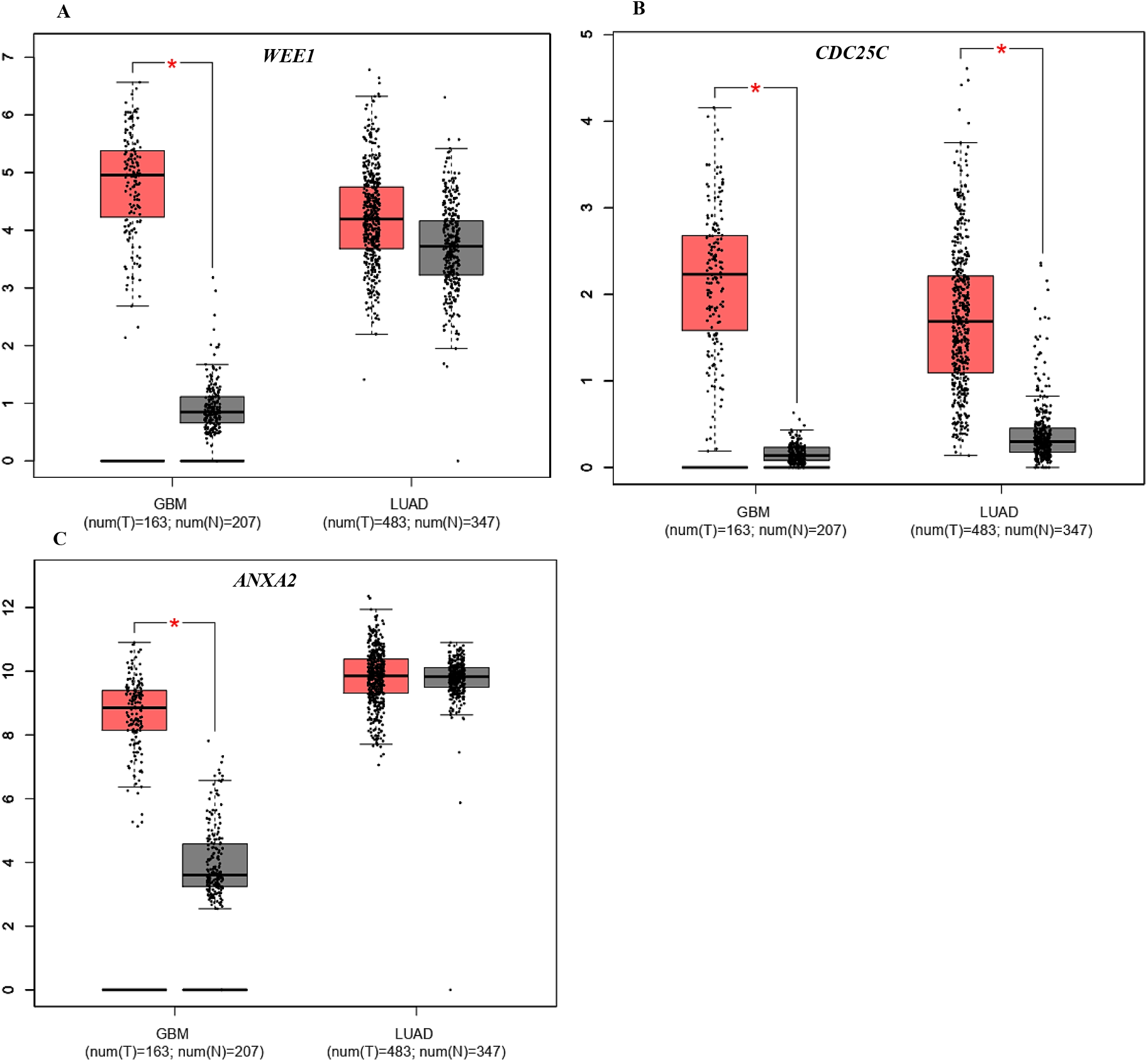
Expression plots of differential expressions of WEE1 (A), CDC25C (B) and ANXA2 (C) in LUAD and GBM compared with respective normal tissues; p-value = 0.01

### 5. Established targets of Withaferin A are also known to be upregulated in Lung Adenocarcinoma and Glioblastoma

Because WA treatment showed marked cell death in GBM cells – A172 and T98G, and lung adenocarcinoma cells – A549, we enquired into role of the validated molecular targets of WA and their relevance in GBM and LUAD. Using GEPIA (Gene Expression Profiling Interactive Analysis), which is a web server that utilizes public RNA sequencing data to perform gene expression analysis, we probed relevance of WEE1, CDC25C and ANXA2, which have been identified as targets of WA [22,23]. Expressions of these genes in TCGA datasets were analyzed for both GBM and LUAD comparing tumor samples to normal tissue against expression levels on log_2_(TPM+1) transformation. While WEE1 and CDC25 were found to be upregulated in both GBM and LUAD, ANXA2 was significantly upregulated only in GBM.

## Discussion

Rapid proliferation of cells comprising a solid tumor creates a hypoxic environment for the cells spatially located within the tumor core. Cancer cells undergo global readaptation of gene-regulatory networks and signalling pathways to thrive under such limiting oxygen and nutrients’ availability. Tumor hypoxia, thus, plays an important role in promoting metastasis, neo-angiogenesis and drug resistance, thus presenting as a negative prognostic factor.

While previous studies have shown WA to have promising pharmacological properties as potential anti-angiogenic, anticancer, antimetastatic and anti-inflammatory; no systematic study of its efficacy in a hypoxic context has been reported in literature to the best of our knowledge [13,24–26]. Based on existing literature, we hypothesized that WA could indeed be a potential candidate small-molecule to target tumor hypoxia and its adaptive consequences. We evaluated its impact in cancer cells of lung and brain growing in conditions of hypoxia *vs*. normoxia. Our study, via cellular and molecular assays, showed that WA is an effective anticancer agent which inhibits both proliferative as well as migratory potential of different cancer cell lines.

In this study, our results clearly indicate that WA is equally effective in inhibiting the proliferative potential of A549 cells, TMZ resistant T98G, A172 and HEPG2 cells in both normoxic as well as hypoxic conditions. Wound healing assay confirmed that WA also curbs migratory potential of A549 cells in an equal measure in both normoxia and hypoxia. The lower concentration (1µM) reduced the migration by 40% and the higher (4µM) curbed it by almost 80%. This equally effective behavior of WA in both normoxia as well as hypoxia uncovers new avenues for exploring its potential in targeting hypoxic tumor microenvironment. This observed efficacy of WA in both normoxia and hypoxia is exciting and the possibility that WA is able to wear down survival advantages provided by HIF dictated transcriptional program appeared worth investigating further.

The detrimental traits that tumor cells acquire as a consequence of hypoxic tumor microenvironment-such as being refractory to drug and radiation therapies, enhanced metastasis, neo-angiogenesis, are a result of transcriptional choreography conducted by the master regulators such as HIF-1α & HIF-2α and NDRG1. WA, in hypoxic conditions, modulates the expression of HIFs as well as its well-known downstream effectors such as NDRG1, GLUT1 and PDK1. Bostrom PJ, et al. (2016), in their study observed that strong expression of GLUT-1 is associated with poor survival in patients with bladder cancer. Together these results indicate that WA may be directly affecting the HIF regulators or at a level upstream of HIF activation [27].

It is a well-established fact that hypoxia in tumors is usually associated with enhanced metastasis via upregulation of LOX, TWIST, Vimentin and N-Cadherin [28]. We show that in concert with cellular assays, in A549 cells, WA treatment downregulated markers of metastasis and EMT such LOX, TWIST, E-Cadherin, N-Cadherin, and Vimentin as assessed by qPCR post 24-hour treatment with WA. Our data indicate that in hypoxic conditions, where LOX expression is more relevant, WA treatment drastically downregulates LOX expression in a dose dependent manner [29]. In 1% hypoxia, doses of 1µM and 4µM downregulated LOX by >8fold and >9fold, respectively. The ability of WA to strongly downregulate LOX, a metastasis enabler, is a highly promising outcome-which can prove to be therapeutically advantageous.

Other inducers of EMT - TWIST, N-Cadherin, Vimentin, are also similarly downregulated on treatment with WA. Expression of Twist and Vimentin is significantly associated with advanced TNM stage. Complementing these results, expression of E-Cadherin was observed to be reduced (> 2.5 folds) in response to 1% hypoxia with respect to normoxia, but is enhanced post treatment with WA. E-Cadherin helps maintain epithelial polarity of cells, keeping invasive potential of cancer cells in check. Previous studies have shown that hypoxia induces EMT to promote invasive and metastatic potential of cancer cells by repressing E-Cadherin, and that expression of E-Cadherin is negatively correlated with expression of Twist [30]. Our results show that repression of inducers of EMT and upregulation of E-Cad are complementary processes and suggest that the cascade of these outcomes might originate either at the repression of hypoxia-inducible factors or upstream of it. However, further studies are required for systematically exploring and validating this.

Reiterating its therapeutic potential, WA treatment, in hypoxic conditions, also increased the expression of pro-apoptotic genes. Our data indicate increased expression of p53, Bax, and p21 in 1% hypoxia post WA treatment. Expression of p53 was induced in untreated hypoxia by almost 10 folds w.r.t untreated normoxia, an observation in sync with other published studies; our treatment with WA (4µM) further enhanced its expression. Studies have shown that severe hypoxia may lead to induction of p53 via HIF-1α, indicating a possible twofold role for HIF-1α, the first being a tendency to promote tumor growth by inducing angiogenesis and the second being that of promoting apoptosis via stabilization of p53 protein [31]. Our observations and results with WA treatment in A549 cells, where p53 expression is observed to increase, indicate that WA might be tilting this balance towards apoptosis, thereby curbing tumor proliferation.

Bioinformatic analysis showed that established targets of WA such as WEE1, ANXA2 and CDC25C are also upregulated in GBM except ANXA2 which is not known to show significant upregulation in LUAD but remarkably high in GBM. ANXA2, a pleotropic gene, is not only known to be involved in cancer and EMT progression but is also hypoxia induced in GBM [32,33]. The exact mechanism how WA targeting ANXA2 in hypoxic context would be leading to cell death as observed in our study will be interesting. Similarly, WEE1 and CDC25C, both playing a crucial role in cell cycle regulation and therefore in progression of various cancers. Studies have shown that inhibiting WEE1 can induce cancer cells to enter mitosis prematurely, leading to mitotic catastrophe and cell death [34]. Inhibitors targeting WEE1 have reached clinical trials underlining the importance of WEE1 as a potential anticancer target [35]. Following this and combining with data obtained from our study, WA also seems to be a potential anticancer agent but effective in tumor hypoxia, too.

This is the first systematic study to explore the effects of WA on cancer cells in the context of hypoxic microenvironment and to report that it is indeed effective in downregulating hypoxia-induced pathways. Also, ours is the first study to show that WA significantly (and reproducibly) inhibits essential and bona fide markers of hypoxia-HIF-1α, HIF-2α and NDRG1. This inhibition of hypoxia regulators is manifested as inhibition of the downstream effectors of the hypoxia-adaptation program such as increased metastasis and invasion. Notably, while TMZ is ineffective in suppressing growth in T98G cells, WA is highly effective in both normoxia as well as in hypoxic conditions. This sensitivity of T98G to WA could most probably be due to the ability of WA downregulate NDRG1, a strong mediator of chemoresistance against TMZ[36]. More studies with T98G and WA need to be performed to further explore this insight.

Collectively, our data indicate that WA is an effective antineoplastic compound not just in normoxic conditions but also in hypoxic microenvironments. The data indicated its efficacy in both non-small cell lung cancer cell lines as well as recalcitrant T98G and A172 glioma cell lines. Combining it with standard of care drugs such as cisplatin in lung cancers and TMZ in glioma shows enhanced efficacy at low dosages. Validating these results in vivo could provide a gateway to address the increased metastasis and drug resistance often associated with hypoxic microenvironments in solid tumors, and would confirm that WA has the potential to serve as an important adjuvant therapy to be used alongside standard treatment regimens.

## Acknowledgement

DM and SG are recipients of Senior Research Fellowship from Council of Scientific and Industrial Research. DM also received Senior Research Fellowship from Indian Council of Medical Research. This project has no direct funding however TS lab is supported by grants from Science and Engineering Research Board, (SERB)-POWER, ICMR-IIRP and DU-Institute of Eminence. We are thankful to Central Instrumentation Facility at UDSC and to DST-FIST and UGC-SAP grants to the Department.

## Notes

### Competing Interest Statement

The authors have declared no competing interest.

