## Supplementary Data for "Withaferin A downregulates NDRG1 to overcome hypoxia mediated EMT and chemoresistance in lung adenocarcinoma and glioma cells"

Supplementary Figures

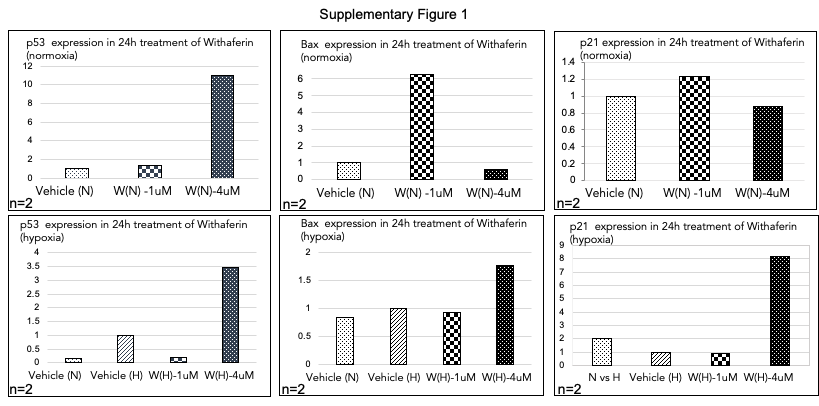

*Supplementary Figure 1 : Fold change values of qRT-PCR results for pro-apoptotic markers.  p53, Bax and p21 expression were studied in A549 lung cancer cells exposed to 24 h treatment of Withaferin A in normoxia and hypoxia. Fold change values were recorded from delta CT calculations.*

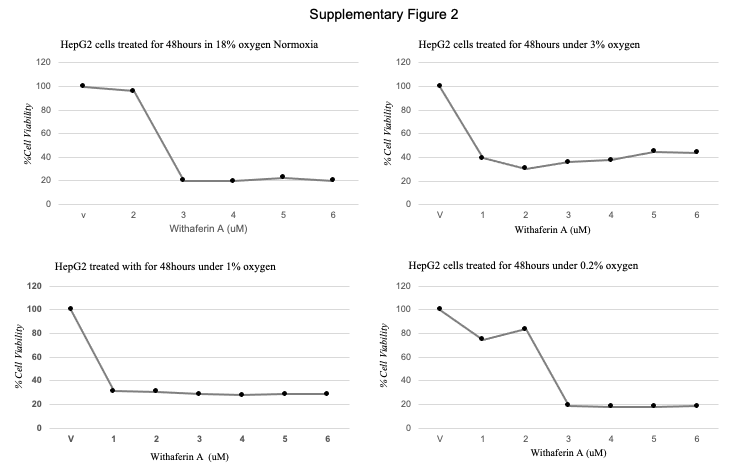

*Supplementary Figure 2: Cytotoxic effect of Withaferin on HEPG2 liver cancer cells in normoxia  and hypoxia (0.2%, 1% and 3%)*

*Table 1 : Primers used in this study*

| *Gene* | *Primer strand* | *Sequence* | *Base pair* |
| --- | --- | --- | --- |
| *B2M* | *Forward* | *GAGGCTATCCAGCGTACTCCA* | *21* |
|  | *Reverse* | *CGGCAGGCATACTCATCTTTT* | *21* |
| *ACTB* | *Forward* | *ATGATGATATCGCCGCGCTC* | *20* |
|  | *Reverse* | *CCACCATCACGCCCTGG* | *17* |
| *18S* | *Forward* | *GTAACCCGTTGAACCCAT* | *18* |
|  | *Reverse* | *CCATCCAATCGGTAGTAGC* | *19* |
| *LOX* | *Forward* | *CGACGACCCTTACAACCCC* | *19* |
|  | *Reverse* | *GTCTGGGAGACCGTACTGGA* | *20* |
| *E-Cadherin* | *Forward* | *GACCCAACCCAAGAATCTATCA* | *22* |
|  | *Reverse* | *AGGCTGTGCCTTCCTACAGAC* | *21* |
| *N-Cadherin* | *Forward* | *ACAATGCCCCTCAAGTGTTAC* | *21* |
|  | *Reverse* | *CATTAAGCCGAGTGATGGTCC* | *21* |
| *GLUT1* | *Forward* | *CAACGCTGTCTTCTATTACTCC* | *20* |
|  | *Reverse* | *CAAACAGCGACACGACAGTG* | *18* |
| *Vimentin* | *Forward* | *CGGGAGAAATTGCAGGAGGA* | *20* |
|  | *Reverse* | *AAGGTCAAGACGTGCCAGAG* | *20* |
| *PDK1* | *Forward* | *TCCTGGACTTCGGATCAGTG* | *20* |
|  | *Reverse* | *TGCAACCATGTTCTTCTAGGC* | *21* |
